# On the interpretation of tractography-based parcellations

**DOI:** 10.1101/409268

**Authors:** Jonathan D. Clayden, David L. Thomas, Alexander Kraskov

## Abstract

Connectivity-based parcellation of subcortical structures using diffusion tractography is now a common paradigm in neuroscience. These analyses often imply voxel-level specificity of connectivity, and the formation of compact, spatially coherent clusters is often taken as strong imaging-based evidence for anatomically distinct subnuclei in an individual. In this study, we demonstrate that internal structure in diffusion anisotropy is not necessary for a plausible parcellation to be obtained, by spatially permuting diffusion parameters within the thalami and repeating the parcellation. Moreover, we show that, in a winner-takes-all paradigm, most voxels receive the same label before and after this shuffling process—a finding that is stable across image acquisitions and tractography algorithms. We therefore suggest that such parcellations should be interpreted with caution.

## Introduction

Diffusion tractography uses proxy information about white matter structure to reconstruct the paths of neural tracts in the living brain. Being based on magnetic resonance imaging (MRI), it is a noninvasive technique with broad applicability in neuroscience and the clinic. An increasingly common application of tractography is connectivity-based parcellation, a paradigm in which a contiguous anatomical region—typically cortical or sub-cortical grey matter—is parcellated into subregions based on the inferred projections from each imaging voxel contained within it. This is established by running diffusion tractography a large number of times and assessing the pattern of connections to a set of target regions.

The steadily growing popularity of the technique has seen it being applied to the parcellation of a wide range of brain structures across many studies. The thalamus was the canonical early example, due to its extensive connectivity to different parts of the cortex, its functional relevance to a range of important neurological disorders and the well-established histological evidence of its nuclear structure (Guillery & Sherman, 2002; Krauth et al., 2010; Morel et al., 1997). Tractography-based *in vivo* thalamic parcellations were demonstrated in a seminal paper by Behrens et al. (2003), and similar principles have since been applied to the amygdala and basal ganglia (Bach et al., 2011; Draganski et al., 2008; Lambert et al., 2012; Saygin et al., 2011). Cortical regions have been explored too: Johansen-Berg et al. (2004) showed a marked distinction between the connectivity of the supplementary motor area (SMA) and the adjoining pre-SMA, Anwander et al. (2007) demonstrated a connectivity-based subdivision of Broca’s area and Jakab et al. (2012) clustered the insula, amongst many other studies.

Use of information from MRI as the basis for such parcellations predates the use of tractography. Magnotta et al. (2000) described a cortex-attenuated MRI sequence able to render thalamic nuclei more visible than on conventional structural scans—although segmentation of the nuclei in such images would require additional processing. Tuch (2002, ch. 6) clustered voxels directly from diffusion MRI data, using a combination of proximity and fibre orientation information. But tractography-based parcellation rapidly superseded such approaches in the literature, and improvements to tractography methods—particularly the ability to resolve fibre crossings within image voxels—led to greater stability in the parcellations (Behrens et al., 2007).

In addition to their value in primary neuroscientific investigations, such parcellations are relevant in the clinic. Patient-specific maps of subcortical nuclei provide potentially valuable navigation information prior to electrode implantation for deep brain stimulation (DBS), a therapeutic neurosurgical intervention used to treat movement disorders such as Parkinson’s disease and dystonia. Recently, da Silva et al. (2017) have demonstrated a connectivity-based parcellation of the globus pallidus internus, a frequent target for DBS, while Akram et al. (2017) identified clusters in the subthalamic nuclei of patients that were associated with alleviation of various Parkinsonian symptoms.

Although the tractography-based approach has been shown to be reproducible, and in many cases to broadly match expectations from neurophysiology and anatomy (Klein et al., 2007), the approach has received some criticism. Eickhoff et al. (2015) reviewed several challenges, including inconsistencies between parcellations derived from different clustering methods and imaging modalities, difficulties with statistical inference in this context, and the role of functional gradients as opposed to clearly defined nuclear boundaries. Nevertheless, the mechanics of the technique are rarely scrutinised in detail. In particular, while tractography itself faces a number of outstanding issues, such as a prepon-derance of “false positive” connections (Maier-Hein et al., 2017), the impact of these and other matters of procedure on the parcellations has not been explored.

In this work we examine the role of fibre orientation information within the parcellated region of interest, by deliberately scrambling the voxels within it and considering the effect on the pattern of projected connections. We demonstrate that coherent structure within the human thalamus in diffusion MRI data is not necessary for plausible delineation of connected subregions. We therefore conclude that such parcellations are potentially prone to overinterpretation, and must be treated with caution.

## Methods

Ethical approval for all imaging was granted by the Research Ethics Committee at University College London.

Diffusion-weighted spin-echo echo-planar (DW-SE-EPI) images were acquired from 17 healthy adult volunteers (six female; mean age at scan 32.84 yr, standard deviation 8.13 yr) on a Siemens Avanto 1.5 T scanner with a 32-channel head coil. Diffusion weighting was applied along 30 noncollinear directions at *b* = 800 s mm^−2^ and 60 directions at *b* = 2400 s mm^−2^, and nine volumes were acquired with *b* = 0. The echo time was 98 ms and the volume repetition time was 8.30 s. 60 slices were acquired, with a matrix size of 96 × 96 and a voxel size of 2.5 mm in each dimension. The data were converted from DICOM to NIfTI-1 format using TractoR (Clayden et al., 2011), corrected for susceptibility and eddy current induced distortions using topup and eddy from FSL version 5.0.11 (Andersson et al., 2003; Andersson & Sotiropoulos, 2016; Smith et al., 2004), and masked using FSL’s brain extraction tool (Smith, 2002). A *T*_1_-weighted structural scan was also acquired (3D Fast Low-Angle SHot; flip angle 15°, echo time 4.94 ms, repetition time 11 ms, resolution 1 × 1 × 1 mm), and automatically parcellated using FreeSurfer (Desikan et al., 2006). The raw data is available online (https://osf.io/94c5t/; Deligianni et al., 2016).

To explore the generalisability of our findings across acquisitions, an additional data set was acquired from a 36 year-old male volunteer on a Siemens Prisma 3 T scanner with 64-channel head coil. In this case, diffusion data were acquired along 60 directions at *b* = 1000 s mm^−2^ and 60 directions at *b* = 2200 s mm^−2^, along with 14 interspersed *b* = 0 volumes. A multi-band DW-SE-EPI sequence was used with multi-band factor 2 (Setsompop et al., 2012); the echo time was 60 ms and the volume repetition time was 3.05 s. 66 slices were acquired with a matrix size of 110 × 110 and a 0.2 mm slice gap; the voxel size was 2 mm in each dimension. The *T*_1_-weighted scan in this case used a Magnetisation-Prepared Rapid Acquisition Gradient Echo sequence with a flip angle of 8°, echo time of 2.74 ms, repetition time of 2300 ms, inversion time of 909 ms and reconstructed resolution of 1 × 1 × 1 mm. All image preprocessing was equivalent.

The left and right thalami, and non-overlapping masks covering the frontal cortex, precentral gyrus, postcentral gyrus, temporal cortex and parietal– occipital cortex, were extracted from the FreeSurfer parcellations (see Fig. 1a). These masks were transformed to diffusion space using NiftyReg (Modat et al., 2010), based on a nonlinear registration between the *T*_1_-weighted image and a *b* = 0 volume from the diffusion acquisition.

**Figure 1:**
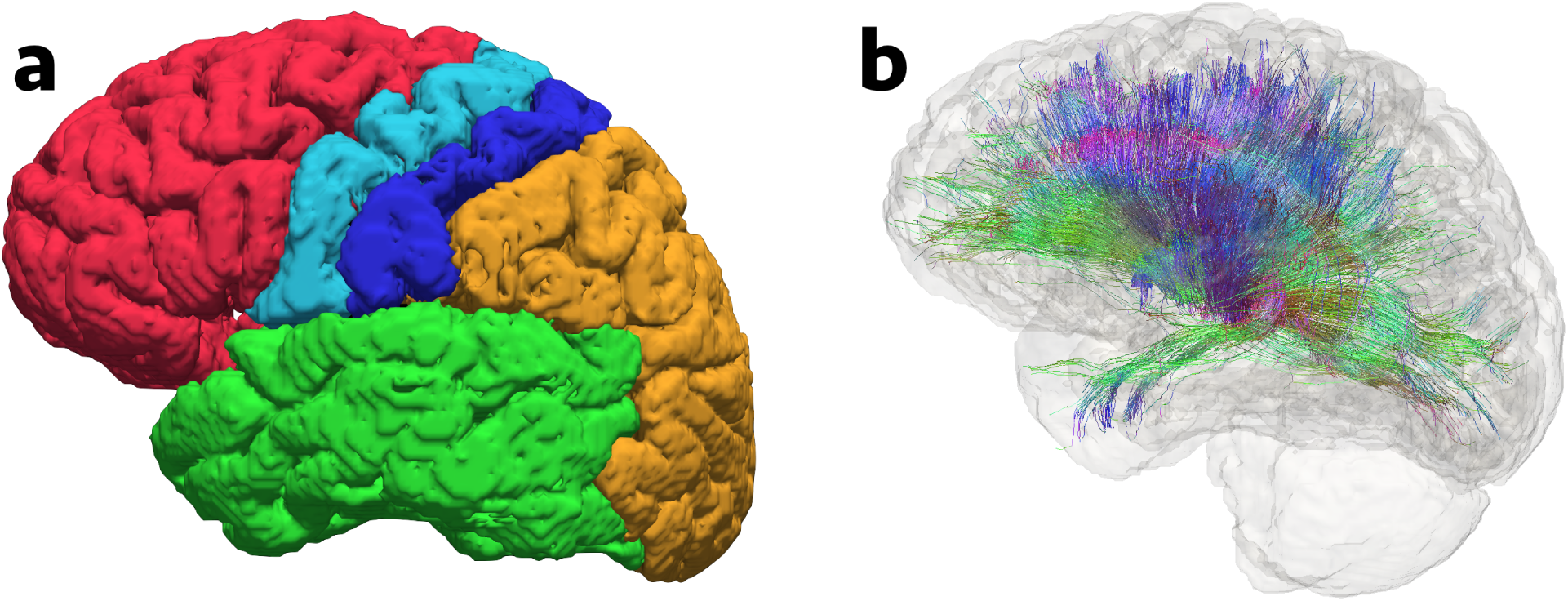
Rendered three-dimensional map of the cortical targets used in this study (a), along with a full set of streamlines projecting from the thalamus (b). Targets are the frontal cortex (red), precentral gyrus (light blue), postcentral gyrus (dark blue), temporal cortex (green) and parietal–occipital cortex (orange). Streamlines are coloured according to their local orientation, using the conventional colour scheme with red for left–right, green for anterior–posterior and blue for superior–inferior.

“Ball-and-sticks” models with one, two and three sticks were fitted separately to each dataset using FSL-BEDPOSTX (Behrens et al., 2007). A copy of each dataset was made, and the fitted ball-and-sticks model parameters for each voxel were randomly repositioned in space within the left and right thalamus in turn (see Fig. 2).

**Figure 2:**
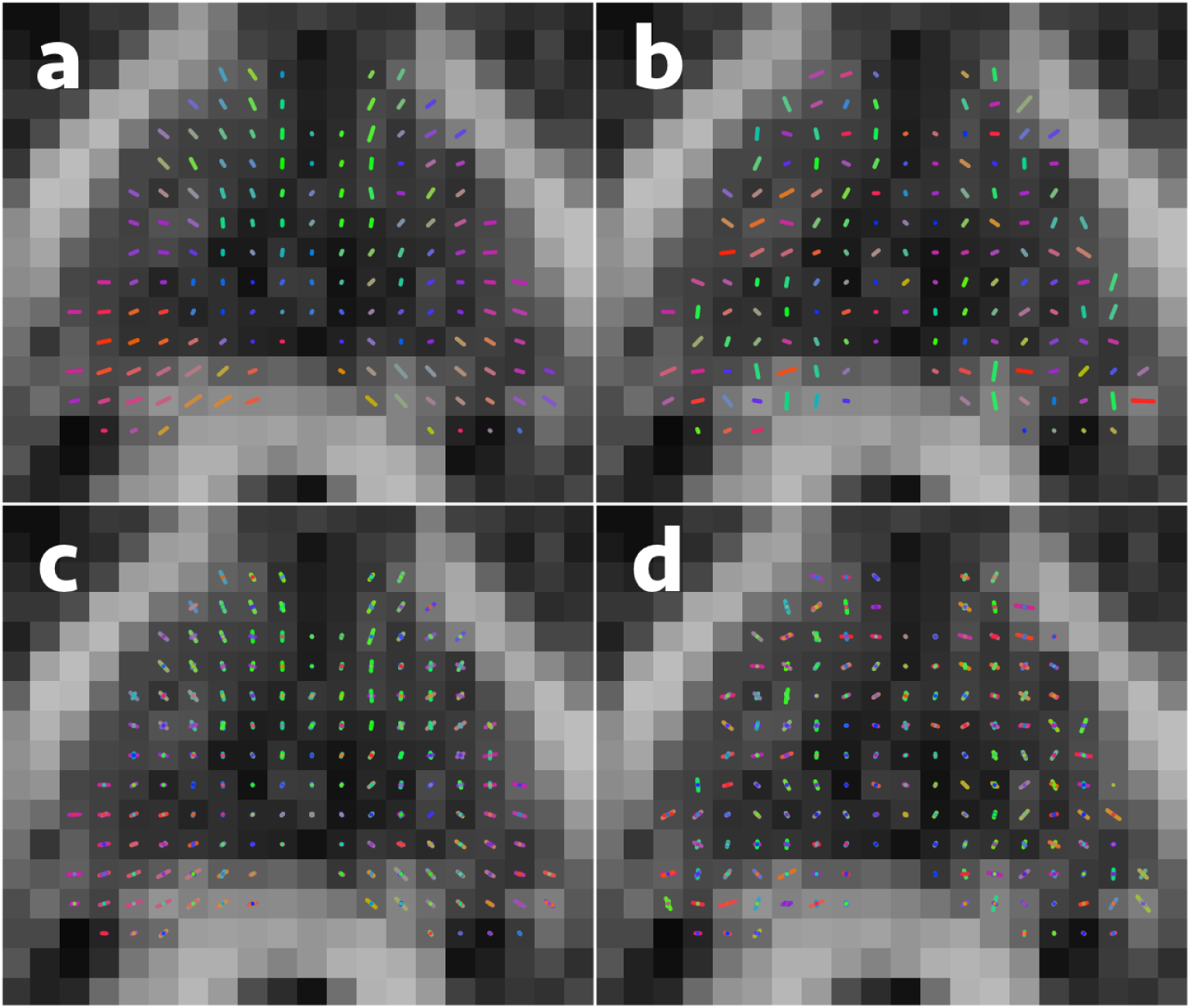
Illustration of the effect of parameter shuffling within the thalamus in one subject. Fibre direction information is shown as fitted by BEDPOSTX for the one stick (a) and three stick (c) cases, alongside the shuffled equivalents (b, d). All parameters are moved wholesale to a randomly chosen other voxel, separately for each hemisphere. The axial slice containing the most thalamus voxels is shown in each case.

In every unshuffled and shuffled dataset, 5000 streamlines were seeded from each voxel within the thalamus masks, and streamlines were propagated using TractoR in one direction only. Streamlines reaching one of the target masks were counted as connecting the source voxel to that target, and then immediately terminated (see Fig. 1b). Each thalamus voxel was coloured to indicate the target at which the greatest number of streamlines arrived, following the “winner-takes-all” scheme introduced by Behrens et al. (2003). No colour was assigned where none of the streamlines from a voxel reached any of the target regions.

Constrained spherical deconvolution (CSD) was also separately performed on the preprocessed 3 T dataset, using MRtrix 3.0 RC3 (Tournier et al., 2007). An iterative algorithm was used to fit the response function (i.e., deconvolution kernel) from the *b* = 2200 s mm^−2^ shell in the diffusion-weighted data, using a maximum spherical harmonic order of eight (Tournier et al., 2013). The default tracking parameters in MRtrix were used to generate 5000 streamlines per seed voxel, except that streamlines were propagated in one direction only, to match the process used with the ball-and-sticks models. Maps of the end-points of streamlines reaching each of the target regions were obtained, and a winner-takes-all parcellation was derived from these maps.

In every case outlined above, a series of summary statistics were compiled, capturing the numbers of streamlines reaching the targets, the numbers of labelled voxels and the degree of consistency in the labels before and after shuffling. To provide a meaningful baseline, the degree of agreement under a direct, random permutation of voxel labels was calculated as described in the Appendix. Analysis was performed with R version 3.5.0 (R Core Team, 2018), and graphics were created with TractoR and the ggplot2 package (Wickham, 2009).

## Results

In every dataset and for every analysis, our shuffling process drastically reduced the number of streamlines reaching any of the target regions, due to the disorderly nature of the orientation information within the thalamus. This resulted in several voxels being unlabelled, especially towards the centre of the thalamus. Nevertheless, the parcellations were still remarkably coherent, with a level of agreement in voxel labels which was well above chance in every case. Switching acquisitions and tractography algorithms did not alter the pattern of the results.

FIG. 3 demonstrates the effect of the shuffling in the three subjects with the greatest, the median and the least amount of agreement in voxel labels before and after shuffling. It shows the axial slice containing the largest number of thalamus voxels in each subject, parcellated as described above, coloured as shown in Fig. 1a and overlaid on a fractional anisotropy map. We observe that parcellations in the unshuffled data generally become a little cleaner as the number of fibre directions modelled per voxel increases (from left to right within each subject)— but the same is also true for the shuffled variants. Although several voxels are uncoloured, clear and coherent structure remains visible in the shuffled parcellations, and there remains evidence of compact nuclei, despite the absence of internal orientational structure within the thalamus in these cases.

**Figure 3:**
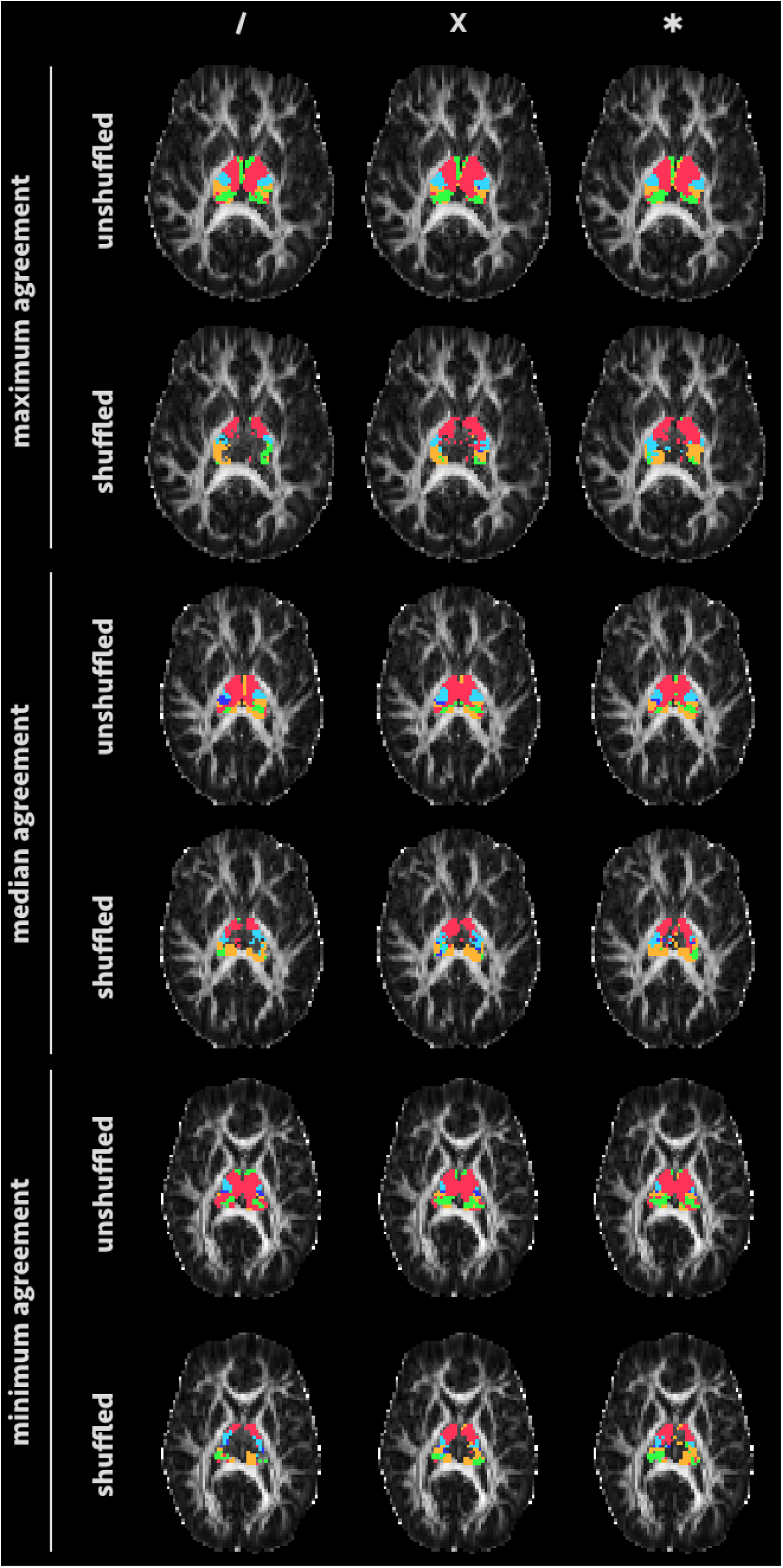
“Winner-takes-all” parcellations before and after spatial shuffling for three subjects, specifically those with the greatest (top), median (middle) and least (bottom) number of voxels sharing a label before and after shuffling. The one-stick case is in the left column, two-stick case in the middle and three-stick case on the right. The axial slice containing the most thalamus voxels is shown in each case, and colours match the cortical areas shown in Fig. 1a.

FIG. 4 shows the distributions of various numerical indicators of the effects of shuffling, across the core dataset of 17 subjects. Averaged across this dataset, just 14–17% as many streamlines seeded from the thalamus reached one of the cortical target after shuffling, compared to before, but 61% (for the one-stick model), 83% (two sticks) and 85% (three sticks) of voxels received a label—compared to almost 100% before shuffling. Out of these connected voxels, 63–66% received the same label (from the five available) after shuffling, compared to a theoretical random baseline averaging 34–35% (cf. the Appendix). Clearly, therefore, the shuffling process has not fully disrupted the parcellation as might be expected.

**Figure 4:**
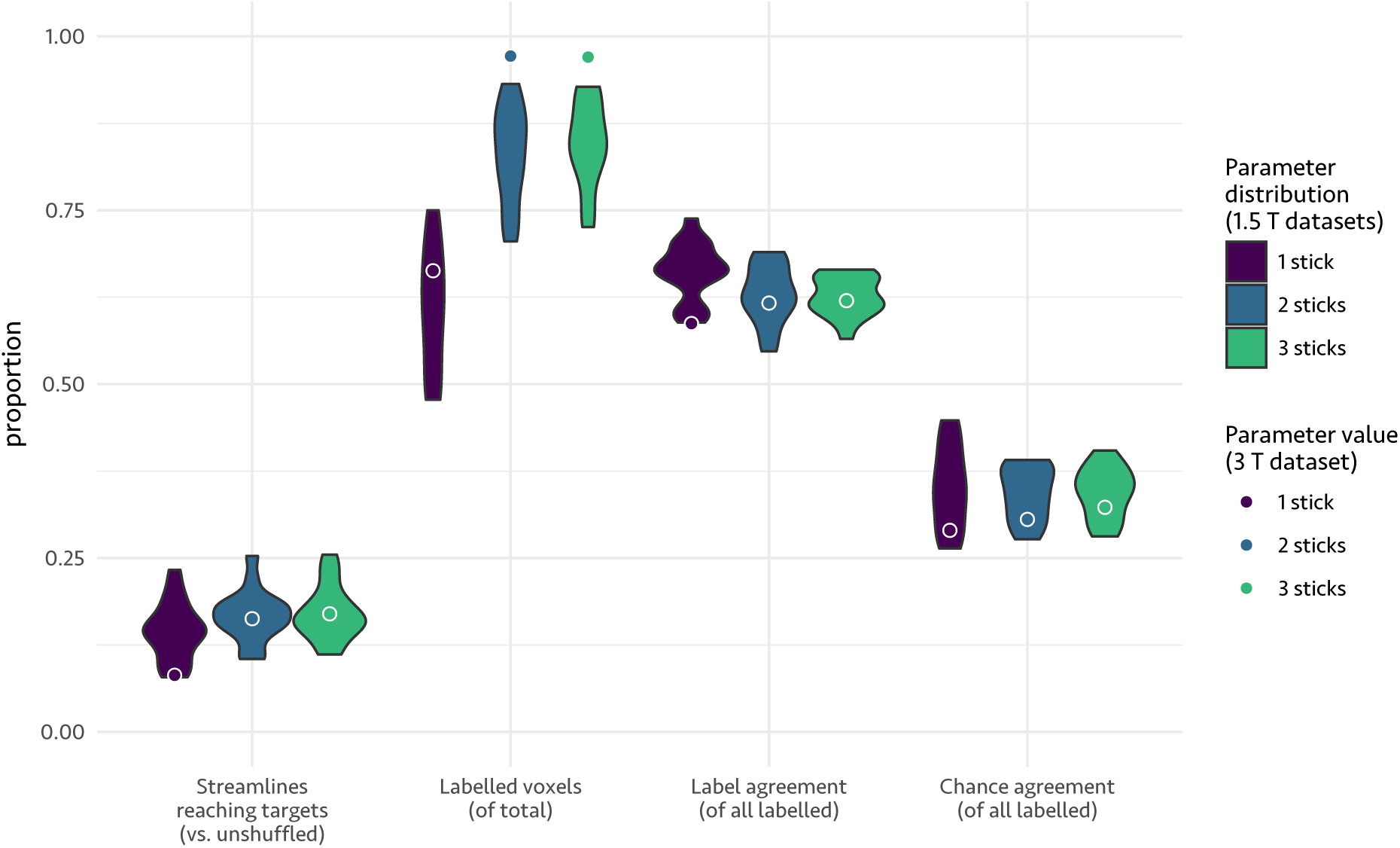
Violin plots quantifying the effects of shuffling across the datasets. Four measures are shown, as proportions: the number of streamlines reaching any of the target regions after shuffling, compared to before; the fraction of voxels receiving a colour; and the proportions of those labelled voxels which receive the same label before and after shuffling, in practice and at chance level. Each violin shows the distribution of values for a particular number of BEDPOSTX stick compartments across the 1.5 T datasets, while the point circled in white shows the position of the 3 T dataset. Note that the chance level varies between datasets, because of the different numbers of thalamic voxels in each case.

The equivalent figures for the single 3 T dataset are shown in Fig. 4 as coloured points circled in white, and the shuffled and unshuffled parcellations are shown in Fig. 5, including for the CSD pipeline. The results are generally similar to those described above, with the numerical values mostly within the range observed in the main dataset. The exceptions are the proportions of labelled voxels in the two and three stick models, which are somewhat higher than in the other cases, perhaps due to the higher image resolution. Nevertheless, the degree of label agreement before and after shuffling is in line with the broader pattern. CSD does not have a fixed bound on the number of fibre populations in the same way as BEDPOSTX, and the numerical results were a little higher throughout, with 25% as many streamlines reaching targets after shuffling, 100% of voxels connected and 75% overlap in labels against a random baseline of 39%. Interestingly, the label overlap between the parcellations derived from CSD and BEDPOSTX (with three sticks) was also 75% (cf. Fig. 5).

**Figure 5:**
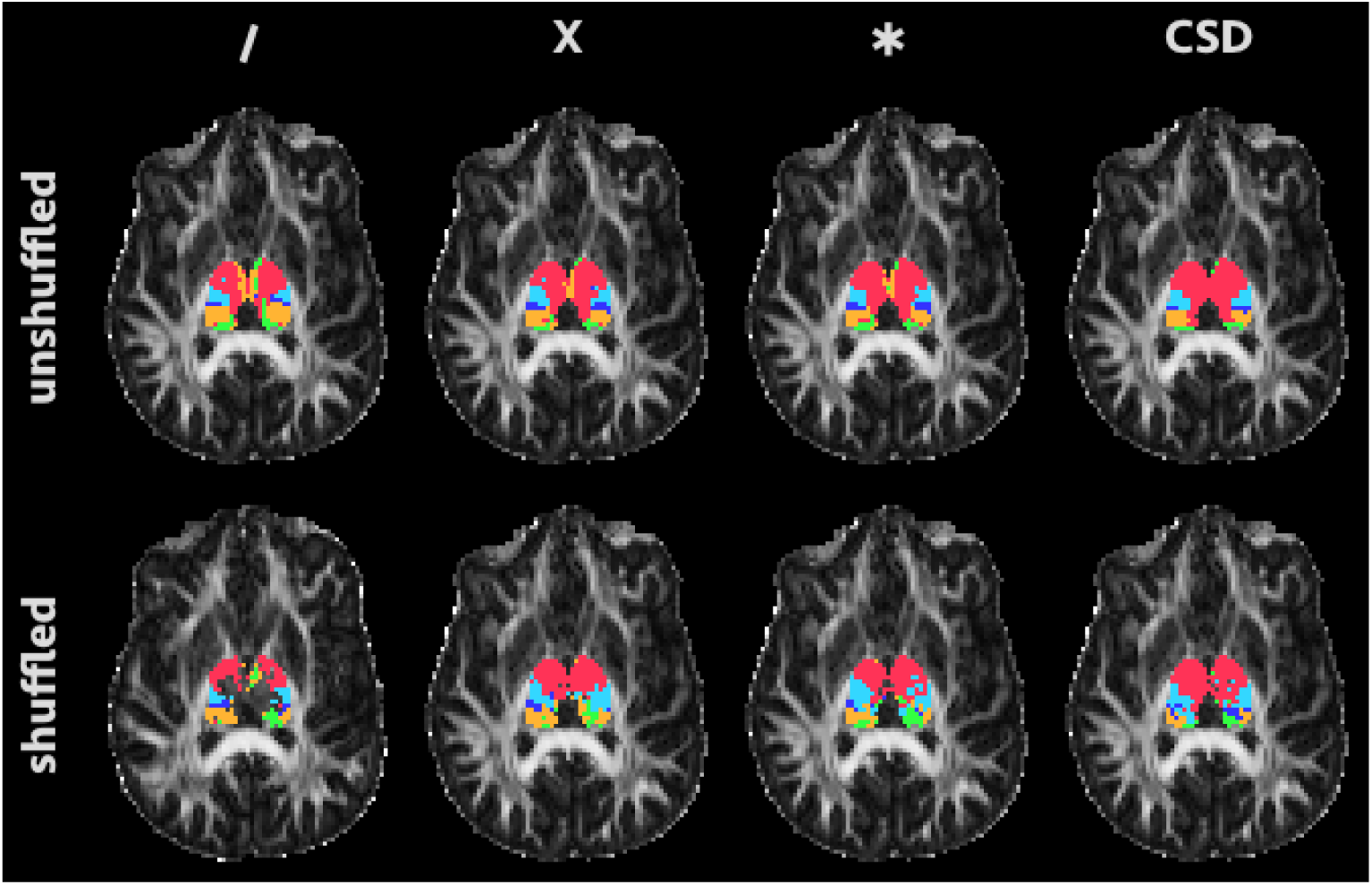
“Winner-takes-all” parcellations in the 3 T dataset, based on ball-and-sticks-based (first three columns) and CSD-based (fourth column) tractography. The axial slice containing the most thalamus voxels is shown in each case, and colours match the cortical areas shown in Fig. 1a.

Finally, Fig. 6 shows variants of the maps from Fig. 3, but focussing on the three-stick model and the subject with the median level of agreement between the labels before and after shuffling. The left column of Fig. 6 shows a variant of the winner-takes-all parcellation where opacity at each voxel is determined by the proportion of connected streamlines that reach the winning region; in other words, it indicates the margin by which the winning target wins that voxel. In practice, it looks very similar to the standard parcellation in Fig. 3, both before and after shuffling. The remaining columns of Fig. 6 show maps pertaining to individual target regions, each of which uses a fixed colour for every voxel, but with opacity normalised across voxels such that the voxel with the greatest number of streamlines reaching that target is opaque. Given that the number of streamlines seeded from each voxel is the same, these maps indicate the relative “strengths” of the connections between individual voxels and the target, and in this case the shuffled maps look quite different, being visibly limited to a few isolated voxels in each case, generally at the boundary with white matter.

**Figure 6:**
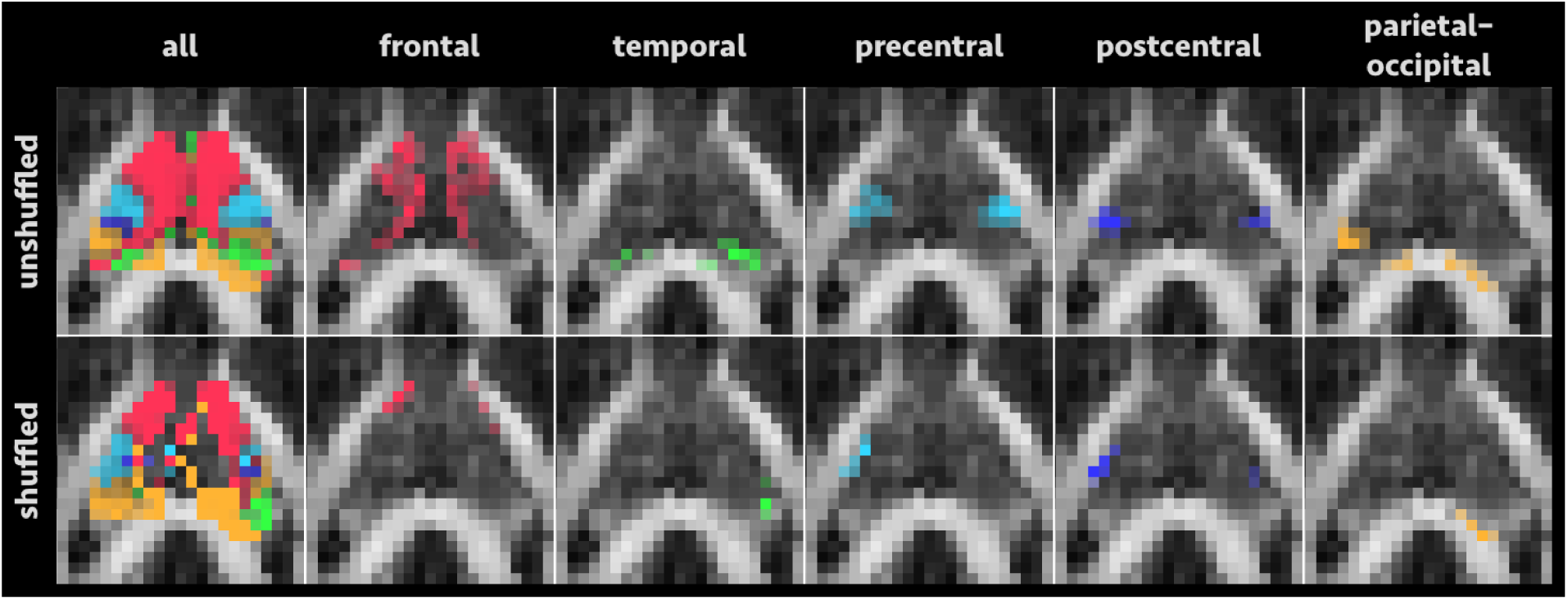
Use of transparency to show the degree of confidence in each voxel label. The first column shows a zoomed-in version of the three-stick subfigure from the median subject in Fig. 3, but with the opacity of each voxel set to indicate the proportion of streamlines from that voxel that reach the winning target region. The remaining columns show maps specific to each target, where the opacity is normalised across voxels such that the voxel with the greatest number of streamlines reaching that target is opaque.

## Discussion

In this study we have demonstrated that artificially destroying internal structure in diffusion anisotropy within the human thalamus does not fully inhibit the creation of plausible streamline-based parcellations, as might be expected. It can be seen directly from Figs 3 and 5 that, while several voxels lose their label completely, the remaining voxels maintain a largely compact, clustered structure, while Fig. 4 shows that consistency in the parcellation before and after shuffling is well above the level expected by chance. Moreover, we have demonstrated that all of these outcomes persist between different image acquisitions and tractography pipelines. Whilst the robustness of these parcellations might be considered to be a positive finding, it casts significant doubt on the implied spatial specificity of voxelwise connectivity-based parcellations, and suggests that plausibility in such parcellations should not be interpreted as strong evidence for a particular subject-specific organisation of subnuclei, or for specific axonal pathways projecting to or from each voxel.

Our shuffling process, illustrated in Fig. 2, is a relatively extreme corruption of the data, which is performed after fitting ball-and-stick model parameters or estimating spherical harmonic coefficients. The advantage of performing this kind of shuffling, rather than using synthetic data, is that all of the parameters are calculated from measurements taken within the thalamus in the same hemisphere, and so they are unquestionably within the realm of general plausibility; they are simply misplaced within the region. Since nothing like this level of corruption would be expected in an unmodified dataset, it is reasonable to expect that very plausible results could be obtained even in the presence of substantial numbers of tractographic false positives and false negatives. Examining the streamline paths themselves, as in Fig. 1b, is highly advisable.

After shuffling, fibre tracking within the thalamus essentially becomes a random walk (cf. Morris et al., 2008), a series of short steps in approximately arbitrary directions. Since tractography algorithms generally impose curvature constraints, this results in a very substantial reduction in the number of streamlines that propagate out of the shuffled region, with the shortest arc lengths being most likely. This explains why most of the unlabelled voxels after shuffling are towards the centre of the thalamus, furthest from the surrounding, unshuffled white matter. Indeed, the relative proximity of each voxel to neighbouring white matter tracts is then the primary determinant of the point of exit from the thalamus, and hence the subsequently inferred connectivity. There are fewer disconnected voxels in models with more complex fibre orientation distributions—two and three stick BEDPOSTX models and CSD—because the fibre tracking process is more likely to be able to identify a viable orientation at each step which does not violate its curvature constraints. In the higher-resolution 3 T dataset, there were fewer disconnected voxels still, perhaps due to the fact that the step distance—which is fixed to 0.5 mm in the TractoR tractography—is larger relative to the voxel size. Finally, the default “iFOD2” algorithm used by MRtrix for tractography is a second-order technique that considers plausible streamline arcs rather than just the immediate local fibre orientation information (Tournier et al., 2010), which may explain why it produces somewhat greater label agreement before and after shuffling.

The consequences of our findings, and the appropriate mitigations for future studies, will depend on the application. A sensible approach in basic neuroscience and clinical studies that treat the parcellation as a primary output would be to seek convergent evidence from other imaging modalities (Glasser et al., 2016; Kelly et al., 2012; Wang et al., 2015). Applying a threshold to the number of connected streamlines needed for a voxel to receive a colour, as demonstrated by Behrens et al. (2003), would also help to avoid tenuous labelling— although such thresholds are inevitably arbitrary. By contrast, in certain clinical contexts it may be that identifying the core locus of connectivity to a particular target region is more important than the parcellation as a whole, and in such cases representations such as those in Fig. 6 would be of more direct relevance, as well as helping to give a greater sense of confidence in the putative subnucleus of interest.

Of course, this study has a number of limitations. We have looked only at the thalamus— although, as we have noted, this is a canonical and well-studied region of interest for parcellation studies. Moreover, due to its relatively large size, issues such as the disconnection of voxels toward the centre of the structure are more visible than might be the case elsewhere. For smaller structures completely surrounded by white matter, such as the subthalamic nucleus, a substantially higher proportion of connected voxels would be expected in the shuffled data, and this may also translate into greater agreement in the parcellation, when compared to the unshuffled original. Additionally, we have considered only the winner-takes-all approach to parcellation, rather than any of the various clustering-based methods. This is partly for simplicity, and partly because winner-takes-all is relatively unambiguous, both methodologically and in terms of its interpretation, whereas more complex multi-target methods (e.g. as used in Draganski et al., 2008) involve more nuance.

In conclusion, we advocate a more cautious interpretation of connectivity-based parcellations than has been prevalent in the literature. We have shown that applying a substantial corruption to diffusion datasets has relatively little effect on such parcellations—indeed, no bigger than switching tractography algorithms. Particularly with modern, multi-compartment diffusion models and tractographic biases such as the tendency to favour directional continuity, algorithms are able to string together paths through even unreliable fibre-orientation fields and brute-force the parcellation. Nevertheless, our findings have no direct bearing on the correctness of the parcellation—indeed, a parcellation of the thalamus with frontal connectivity towards the anterior margin and occipital connectivity posterior is logical from the point of view of economical wiring in the brain (Laughlin & Sejnowski, 2003; Van Essen, 1997). But the lack of sensitivity to internal structure undermines the principle of mapping spatially specific, voxel-to-region connectivity *in vivo*, which is the technique’s core idea.

## Acknowledgments

The authors are grateful to Dr Kiran Seunarine and Prof. Chris Clark for the 3 T volunteer dataset used in this study, and to Prof. Daniel Alexander and Alina Matis for initial discussions about this work. DLT is supported by the UCL Leonard Wolfson Experimental Neurology Centre (PR/ylr/18575). AK is supported by the Wellcome Trust.

## Appendix: Random shuffling

Any given classification of *N* voxels separates them into *n* groups containing *m*_*i*_ elements each, with *i* ∈ {1, 2,…, *n*}, such that 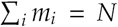. A comparison of the overlap between two such classifications should consider as a baseline the degree of overlap that would be expected by chance.

The number of possible permutations of the labels is given by the multinomial coefficient,

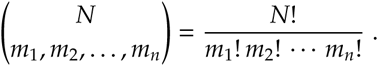

After a random permutation of the labels, the number of cases in which a voxel from group *i* retains that label is

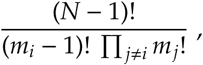

the number of combinations of all elements except the one staying the same.

Taking *X*_*l*_, with *l* ∈ {1, 2, …, *N*}, to be the random variable with value 1 if voxel *l* from group *i* retains its label, and 0 otherwise, we divide the two previous equations to obtain

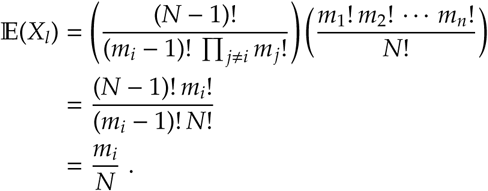

Taking *X* = *X*_1_ + *X*_2_ + *…* + *X*_*N*_, we can make use of the linearity of expectations to quickly determine that

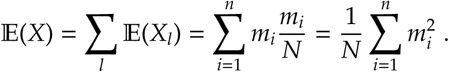

This process is a permutation, and therefore assumes that the proportions of each label remain exactly equal. However, if instead we draw voxel labels at random such that the probability of each voxel having label *i* is *m*_*i*_*/N*, independently of all other voxels, the expectation does not change. Clearly, Pr(*X*_*l*_ = 1) = *m*_*i*_*/N*, since voxel *l* originally had label *i*, and so we can proceed using linearity of expectations as above. Assuming *fixed probabilities* therefore has the same expected outcome as assuming *fixed proportions*.

